## Supplemental tables for "Programmable epigenome editing by transient delivery of CRISPR epigenome editor ribonucleoproteins"

### Supplementary data

| Name | Source | Identifier |
|---|---|---|
| pBS-CMV-gagpol | Addgene | #35614 |
| pCMV-MMLVgag-3xNES-ABE8e | Addgene | #181751 |
| pCMV-VSV-G | Addgene | #8454 |
| CRISPRoff-v2.1 | Addgene | #167981 |
| TETv4 | Addgene | #167983 |
| pU6-sgRNA-lentiviral | Addgene | # 217306 |
| pU6-sgRNA-transient | This study | TBA |
| pCMV-MMLVgag-dCas9 | This study | TBA |
| pCMV-MMLVgag-dCas9-ZIM3 | This study | TBA |
| pCMV-MMLVgag-dCas9-KOX1 | This study | TBA |
| pCMV-MMLVgag-D3A-3L-dCas9 | This study | TBA |
| pCMV-MMLVgag-D3A-3L-dCas9-ZIM3 | This study | TBA |
| pCMV-MMLVgag-D3A-3L-dCas9-KOX1 | This study | TBA |

**Supplementary Table 1. Recombinant DNA.**

| Name | Target | Protospacer sequence (5'-3') |
|---|---|---|
| mU6-CLTA_CRi_gRNA | *CLTA* | CTCCCAGTCGGCACCACAG |
| mU6-H2B_CRi_gRNA | *H2B* | GTAAGACACAGTACAAACG |
| mU6-CD29_CRi_gRNAa | *CD29* | AGAGGAATGCGTTTCCGGA |
| mU6-CD29_CRi_gRNAb | *CD29* | CCGGGCCGGGCTGACGCGG |
| mU6-CD29_CRi_gRNAc | *CD29* | AGAGGCCCAGCGGGAGTCG |
| mU6-CD55_CRi_gRNAa | *CD55* | GCTGGGCGTAGCTGCGACT |
| mU6-CD55_CRi_gRNAb | *CD55* | GGCGCGCCGGGTTAGAACA |
| mU6-CD55_CRi_gRNAc | *CD55* | CTGCGACTCGGCGGAGTCC |
| mU6-CD81_CRi_gRNAa | *CD81* | GAGAGCGAGCGCGCAACGG |
| mU6-CD81_CRi_gRNAb | *CD81* | GGCCTGGCAGGATGCGCGG |
| mU6-CD81_CRi_gRNAc | *CD81* | CCGAAACGCGCCAAGTTGG |
| mU6-CD151_CRi_gRNAa | *CD151* | CCGGACTCGGACGCGTGGT |
| mU6-CD151_CRi_gRNAb | *CD151* | TGTCCAGGGACAATGAGCA |
| mU6-CD151_CRi_gRNAc | *CD151* | GACAATGAGCAGGGTGTCC |
| mU6-MAPT_CRi_gRNAa | *MAPT* | AGTCGACAGAGGCGAGGAC |
| mU6-MAPT_CRi_gRNAb | *MAPT* | GCGGCGCTGCTGTTGGTGC |
| mU6-MAPT_CRi_gRNAc | *MAPT* | TGTTGGTGCCGGAGCTGGT |
| mU6-MAPT_CRi_gRNAd | *MAPT* | CTGCTGTTGGTGCCGGAGC |
| mU6-GAL4_NT_gRNA | *GAL4* | ACGACTAGTTAGGCGTGTA |

**Supplementary Table 2. sgRNA sequences.**

| Antibody or dye | Supplier | Clone | Identifier | Dilution |
|---|---|---|---|---|
| Anti-human CD29-APC | Biolegend | TS2/16 | #303008 | 1:500 |
| Anti-human CD55-APC | Biolegend | JS11 | #311312 | 1:500 |
| Anti-human CD55-FITC | Biolegend | JS11 | #311306 | 1:500 |
| Anti-human CD81-PE | Biolegend | 5A6 | #349506 | 1:250 |
| Anti-human CD81-PB | Biolegend | 5A6 | #349516 | 1:500 |
| Anti-human CD151-PE | Biolegend | 50-6 | #350408 | 1:500 |
| Zombie Aqua Viability Dye | Biolegend | - | #423101 | 1:1000 |

**Supplementary Table 3. Flow cytometry antibodies and dyes.**
